# Effects of high pCO_2_ on snow crab larvae: Carryover effects from embryogenesis and oogenesis reduce direct effects on larval survival

**DOI:** 10.1101/2022.10.06.511100

**Authors:** W. Christopher Long, Katherine M. Swiney, Robert J. Foy

**Author notes:** Corresponding author, (WCL).

## Abstract

Ocean acidification, a decrease in ocean pH with increasing anthropogenic CO_2_ concentrations, is expected to affect many marine animals. We determined the effects of ocean acidification on the economically important snow crab, *Chionoecetes opilio*. By holding females in treatment pH for two brooding cycles and using the resulting larvae, we assessed carryover effects from oogenesis and embryogenesis. Ovigerous females were held at three pHs: ~8.1 (Ambient), 7.8, and 7.5. Larvae were exposed to the same pH treatments in a fully crossed experimental design. Starvation-survival, morphology, condition, and calcium/magnesium content were assessed for larvae. In the first year, starvation-survival of larvae reared at ambient pH but hatched from embryos reared at reduced pH was lowered; however, the negative effect was eliminated when the larvae were reared at reduced pH. In the second year, there was no direct effect of either embryo or larval pH treatment, but larvae reared as embryos at reduced pH survived longer if reared at reduced pH. Larvae hatched from embryos held at pH 7.5 had lower calcium content right after hatching, but the effect was transitory in the second year. There was no effect of larval treatment on calcium content or effect of embryo or larval treatment on magnesium content. Larval morphometrics were slightly altered, though effect sizes were small smaller in the second year. These results suggest both that larvae are highly tolerant of reduced pH, and that embryos are able to acclimate to low pH and this effect carries over to the larval stage.

## Introduction

The anthropogenically-driven increase in atmospheric CO_2_ since the beginning of the industrial revolution has resulted in commensurate increases in the pCO_2_ in the world’s oceans [1]. This causes a host of changes to oceanic carbonate chemistry including a lowering of the pH and the saturation state of calcium carbonate with the average surface pH of the global ocean dropping by about 0.1 pH units since the late 19^th^ century with further reductions predicted as CO_2_ levels continue to rise [1–3]; these changes are collectively referred to as ocean acidification (OA). These changes to oceanic chemistry are known to have physiological effect on many marine organisms, although the effects are highly variable both within and among species, and there is concern that OA could cause changes to marine ecosystems [4,5].

Decapod crustaceans are an ecologically and economically important taxa that can be affected by OA, although effects are highly variable. When exposed to reduced pH many species are able to buffer their hemolymph to achieve acid-base homeostasis, although this is not effective in some species. Buffering is primarily achieved through active ion-transport in the gills, either by Na^+^/H^+^ or Cl^-^/HCO_3_^-^ exchange [6]; however, this can be energetically expensive. Prolonged exposure to OA can result in a range of negative effects including decreased growth [7,8], increased mortality [9,10], altered shell structure and strength [11,12], deformities [13,14], altered behavior [15,16], and decreased fecundity [17], although many species appear to be unaffected [18–21]. For species that are negative affected, OA is projected to result in decreased abundance which, for commercially important species, translates into reduced catches and profitability in the fisheries that depend on them.

In addition to direct effects, OA can have carryover effects where exposure to low pH at one life history stage affects the fitness of later stages [22]. Sometimes exposure at one stage decreases the vulnerability of later life history stages to OA, presumably through phenotypic plasticity [23]. In other cases, exposure at one stage can have continued negative effects on later stages even if those later stages are not exposed to OA [24,25]. Maternal effects, a subtype of intergenerational effects, are another type of carryover effect in which exposure of the mother to OA prior to reproduction affects the fitness of her offspring. Again the effects can be positive or negative [23,26]. Quantifying carryover effects is essential because they are capable of either magnifying or reducing the overall effects of OA on an individual and thus substantially effecting both the magnitude and directionality of overall effects [22].

Snow crabs, *Chionoecetes opilio*, are an important fisheries species in the Bering Sea as well as in the Arctic and north Pacific and Atlantic Oceans. In the Bering Sea, larval crab hatch out in the late spring and early summer [27] and spend about 3-4 month in the plankton before settling to the benthos and molting to the first crab stage [28]. It takes about 4.5-7.5 years for female crabs to reach sexual maturity; females have a terminal molt to maturity that occurs typically in the late winter after which they mate and extrude their first clutch of eggs [27]. In the Bering Sea, embryos are typically brooded for about 15 months for primiparous females or 12 months for multiparous; however, in cold enough waters the females switch to a biennial cycle [29–32].

Nothing is known about the effects of OA on snow crab. However, a closely related species, Tanner crab, *Chionoecetes bairdi*, is strongly affected. Although embryos are not directly affected, maternal effects are strongly negative such that exposure of the mother to OA during oogenesis reduces hatching success by more than 70% [17]. Similarly, although direct effects on the larvae are negligible, juvenile Tanner crabs exhibit decreased growth, increased mortality, and decreased calcification when exposed to OA. Larval Tanner crab are relatively resilient to OA; however, there are also negative carryover effects from exposure during embryo development and during oocyte development [33]. Juvenile Tanner crabs likewise have higher mortality, lower survival, and lower calcification under OA conditions [34]. Finally, in mature female Tanner crab, OA increases hemocyte mortality and decreases intracellular pH [35] and causes internal and external erosion of the exoskeleton which substantially weakens it [12]. We hypothesized that as snow crab are so closely related to Tanner crab that they were likely to be similarly vulnerable to OA. This paper describes a study that examines the direct and carryover effects of OA on snow crab larvae.

## Methods

### Overview

This paper describes a series of experiments on the effect of OA on snow crab larvae that took place over two years. Female snow crab with newly extruded uneyed eggs were captured on the 2014 eastern Bering Sea trawl survey [36] and transported to the Kodiak Fisheries Research Center. They were held at three treatment pHs (see below) throughout brooding. This paper describes a series of experiments that were performed on the larvae in a design that fully crossed embryo pH treatment with larval pH treatment. This allowed us to determine the direct effects of OA on the larvae and the carryover effects from embryogenesis. The females were then provided with a mate in the laboratory (captured at the same time as the females), extruded a second batch of eggs, and kept in their treatment water for a second year. The larvae that hatched were then used in a second series of experiments identical to the first year. This allowed us to determine both the direct effects of OA on the larvae and the carryover effects from both oogenesis and embryogenesis combined.

### Water Acidification and experimental conditions

Both the females and the larvae were exposed to the same three pH treatments based on projected future ocean pH levels: current surface ambient (~pH 8.1), pH 7.8 (projected for 2100), and pH 7.5 (projected for 2200). Sand filtered seawater from Trident Basin intakes at 15 and 26m was used throughout this experiment. Female holding tanks (one per pH treatment) were kept at 2 ± 0.5 °C by recirculating chillers. Females were fed to excess twice a week on a diet of chopped frozen squid and herring. Acidified water for both the female holding tanks and the larvae experimental tanks was produced as detailed in Long et al. [34]. In brief, water was acidified down to ~pH 5.5 by bubbling CO_2_. This water was mixed with seawater in head tanks using peristaltic pumps controlled Honeywell controllers and Durafet III pH probe in the head tanks down to the appropriate pH or the head tanks were controlled via a Durafet III pH probe and direct bubbling of CO_2_. Water from the head tanks then flowed into larvae experimental tanks which were kept at 3 ± 0.5 °C by recirculating chillers. Temperature and pH (total scale) were measured in experimental units daily using a Durafet III pH probe calibrated with TRIS buffer [37]. Water from the head tanks was sampled once per week and samples were poisoned with mercuric chloride and analyzed for dissolved inorganic carbon (DIC) and total alkalinity (TA) at an analytical laboratory. DIC and TA were determined using a VINDTA 3C (Marianda, Kiel, Germany) coupled with a 5012 Coulometer (UIC Inc.) according to the procedure in [38] using Certified Reference Material from the Dickson Laboratory (Scripps Institute, San Diego, CA, USA; [39]). The other components of the carbonate system were calculated in R (V3.6.1, Vienna, Austria using the seacarb package [40].

The treatment pH levels in the two reduced pH treatments averaged the target levels exactly in both years and the ambient treatment averages were only 0.02 pH units different between the two years (Table 1). The ambient pH treatment was supersaturated with regards to both calcite and aragonite while the pH 7.8 treatment was undersaturated with regards to aragonite and the pH 7.5 treatment was undersaturated with regards to both (Table 1). Female holding PH_F_S averaged 8.11 ± 0.08, 7.80 ± 0.02, and 7.50 ± 0.02 in the ambient, pH 7.8, and pH 7.5 treatment respectively and other water chemical properties in the female holding tanks were very similar to those of the larvae [41].

**Table 1.** Water physical and chemical parameters in two years of larval experiment.

| Treatment | Temperature<br>°C | pH<sub>F</sub> | pCO<sub>2</sub><br>μatm | HCO<sub>3</sub><sup>-</sup><br>mmol/kg | CO<sub>3</sub><sup>-2</sup><br>mmol/kg | DIC<br>mmol/kg | Alkalinity<br>mmol/kg | Ω<sub>Aragonite</sub> | Ω<sub>Calcite</sub> |
| --- | --- | --- | --- | --- | --- | --- | --- | --- | --- |
| **2015** |  |  |  |  |  |  |  |  |  |
| Ambient | 3.40 ± 0.17 | 8.16 ± 0.04 | 314.79 ± 31.00 | 1.86 ± 0.02 | 0.10 ± 0.01 | 1.98 ± 0.01 | 2.12 ± 0.01 | 1.56 ± 0.13 | 2.49 ± 0.21 |
| pH 7.8 | 3.17 ± 0.15 | 7.80 ± 0.02 | 751.58 ± 24.49 | 1.99 ± 0.01 | 0.05 ± 0.00 | 2.08 ± 0.01 | 2.12 ± 0.01 | 0.74 ± 0.02 | 1.18 ± 0.03 |
| pH 7.5 | 3.36 ± 0.21 | 7.50 ± 0.02 | 1526.01 ± 54.95 | 2.05 ± 0.01 | 0.03 ± 0.00 | 2.16 ± 0.01 | 2.12 ± 0.00 | 0.39 ± 0.02 | 0.62 ± 0.03 |
| **2016** |  |  |  |  |  |  |  |  |  |
| Ambient | 2.77 ± 0.16 | 8.14 ± 0.03 | 322.23 ± 32.83 | 1.86 ± 0.02 | 0.10 ± 0.01 | 1.98 ± 0.01 | 2.12 ± 0.01 | 1.48 ± 0.12 | 2.37 ± 0.19 |
| pH 7.8 | 2.59 ± 0.17 | 7.80 ± 0.02 | 768.68 ± 24.26 | 1.99 ± 0.00 | 0.05 ± 0.00 | 2.08 ± 0.00 | 2.13 ± 0.01 | 0.69 ± 0.02 | 1.11 ± 0.03 |
| pH 7.5 | 2.75 ± 0.17 | 7.50 ± 0.02 | 1556.47 ± 59.39 | 2.02 ± 0.02 | 0.02 ± 0.00 | 2.14 ± 0.02 | 2.12 ± 0.01 | 0.36 ± 0.02 | 0.58 ± 0.02 |

Water parameters in experimental inserts during larval snow crab experiments. Temperature and pH were measured daily, dissolved
inorganic carbon (DIC) and alkalinity were measured weekly, and all other parameters were calculated. Values are mean ± 1 SD.

### Larval experiments

Larval experiments were identical to those in Long et al. [33], except that the water temperature was kept lower during these experiments because snow crab have a lower temperature range than Tanner crab (*Chionoecetes baridi*). Larval experiments were conducted in inserts made from cut pieces of PVC piping with mesh bottoms placed inside larger treatment tanks. Each insert received water flow of 100 ml/min via a recirculating submersible pump. Survival experiments were performed in inserts with a volume of ~1 L and all other experiments in inserts with a volume of ~2 L. Larvae were unfed during all experiments. During hatching females were held in individual containers and the number of larvae hatched each day was estimated. Larvae experiments were started once enough females within a treatment were at or near peak hatch. Newly hatched larvae for use in the experiments were collected overnight and pooled from 3 females per treatment per year; because females from the different treatments were not all synchronous in their hatching each year, there was some small variation among embryo treatments in the start date for the different experiments. For each experiment embryo treatment was fully crossed with larval treatment for a total of 9 treatments. Five replicates (inserts) were run within each of those 9 treatments for a total sample size of 45 for each experiment.

For starvation survival experiments, 20 larvae were counted out and placed in inserts. They were monitored daily and dead larvae were removed. Each trial was ended when all larvae had died or after 7 weeks had passed (no trial had more than 1 surviving larvae in it when the experiment ended). Mortality data were fit to a series of models using maximum likelihood and assuming a binomial distribution of data. Mortality was modeled with a power formulation of the logistic regression:

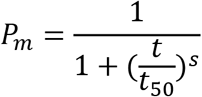

Where *P_m_* is the probability of mortality at time *t*, *t_50_* is the time to 50% mortality (also known as the LT_50_) and *s* is a slope parameter. A series of models were developed in which *t_50_* and s were allowed to vary linearly with embryo treatment, larval treatment and their interaction. The Akieke’s Information Criterion corrected for small sample size (AICc) was calculated for each model and the most parsimonious model selected. Models whose AICcs differed by less than two were considered to have equal weight [42]. Data from each year was analyzed separately.

Condition experiments and calcification experiments were performed in separate trials but using similar methods. When beginning each experiment, samples from the initial pooled larvae used in the experiments were taken and analyzed for dry mass, carbon and nitrogen (CN) content, and calcium and magnesium content as described below. For both experiments, ~300 larvae (estimated volumetrically) were transferred into each insert. Larvae were checked daily and dead larvae were removed by siphoning them off the bottom. The larvae were exposed to treatment water for 7 days and then trials were ended. From the condition experiment, 50 larvae per replicate were counted out, sieved, rinsed briefly in fresh water, and dried. The mass was determined and the average larval dry mass calculated. The remaining larvae in each insert were sieved, rinsed and dried and the CN contents were determined (the 50 larval used to determine the dry mass were recombined with the remaining larvae from each insert to ensure an adequate sample mass). The carbon and nitrogen content was determined at the University of Santa Barbara using the Dumas combustion method and an automated organic elemental analyzer [43]. For the calcification experiment, the larvae from each replicate were sieved, rinsed, and dried and analyzed for calcium and magnesium content at Gel Laboratories via inductively coupled plasm-atomic emission spectrometry. Larval dry mass, carbon, nitrogen, calcium, and magnesium contents, and the C:N ratios were analyzed with fully crossed 2-way ANOVAs with Embryo and Larval treatments as the factors for each year. Where differences were significant Tukey’s post hoc test was used to determine differences among treatments.

Larval morphometry was determined by measuring 15 larvae from the larvae pooled for the condition experiment from each treatment from each year. Digital micrographs were taken of each larvae using a stereo microscope and six measurements were made on each larvae using Image-Pro Plus v. 6.00.260 (Media Cybernetics, Inc., Bethesda, MD, USA): carapace width (including spines), lateral spine length, dorsal spine length, rostro-dorsal length, rostral spine length, and protopodite length as per Long et al. [33]. Measurements were normalized (expressed in terms of the z-scores) before analysis. Morphometry was analyzed using an analysis of similarity (ANOSIM) and visualized using a non-metric multidimensional scaling both performed on a similarity matrix using Euclidean distance. Where there were significant differences among treatments a Similarity Percentage (Simper) analysis was performed on the raw (non-normalized) measurements to determine which measurements most strongly influenced the differences among treatments.

## Results

All raw data from these experiments and the associated metadata are available at the National Centers for Environmental Information [44].

In both years the best fit model of starvation survival was one where both the *t_50_* and *s* parameters varied with both embryo and larval pH treatments and their interaction (Table 2). In the first year, embryo treatment had a larger effect than larval treatment with larvae that were in ambient water as embryos surviving longer than those that were in pH 7.8 or 7.5 water; however larvae that were held in acidified water as embryos survived longer when exposed to acidified water as larvae than when held in ambient water (Fig. 1, Table 3). In the second year, embryo treatment had almost no effect on larvae that were subsequently held in ambient water as larvae; however, larvae from embryos held in pH 7.8 and 7.5 water survived longer than their Ambient counterparts when held in acidified water with larvae surviving longest when exposed to the same pH water as larvae as they were when embryos (Fig. 1, Table 3).

**Table 2:** Ranking of snow crab larvae starvation survival models.

| Model | K | Year 1 |  |  |  | Year 2 |  |  |  |
| --- | --- | --- | --- | --- | --- | --- | --- | --- | --- |
|  |  | AIC<sub>c</sub> | ΔAIC<sub>c</sub> | Likelihood | AIC<sub>c</sub> Weight | AIC<sub>c</sub> | ΔAIC<sub>c</sub> | Likelihood | AIC<sub>c</sub> Weight |
| $t_{50}, s$ | 2 | 3523.66 | 476.79 | 0.00 | 0.00 | 3943.76 | 235.03 | 0.00 | 0.00 |
| $t_{50}(E), s$ | 4 | 3344.38 | 297.51 | 0.00 | 0.00 | 3899.79 | 191.06 | 0.00 | 0.00 |
| $t_{50}(L), s$ | 4 | 3381.45 | 334.58 | 0.00 | 0.00 | 3947.29 | 238.56 | 0.00 | 0.00 |
| $t_{50}(E,L), s$ | 6 | 3158.61 | 111.74 | 0.00 | 0.00 | 3903.04 | 194.31 | 0.00 | 0.00 |
| $t_{50}(E\#L), s$ | 10 | 3120.76 | 73.89 | 0.00 | 0.00 | 3850.76 | 142.03 | 0.00 | 0.00 |
| $t_{50}, s(E)$ | 4 | 3448.34 | 401.47 | 0.00 | 0.00 | 3922.66 | 213.92 | 0.00 | 0.00 |
| $t_{50}, s(L)$ | 4 | 3502.29 | 455.41 | 0.00 | 0.00 | 3861.80 | 153.07 | 0.00 | 0.00 |
| $t_{50}, s(E,L)$ | 6 | 3416.76 | 369.89 | 0.00 | 0.00 | 3847.21 | 138.48 | 0.00 | 0.00 |
| $t_{50}, s(E\#L)$ | 10 | 3401.75 | 354.88 | 0.00 | 0.00 | 3803.82 | 95.09 | 0.00 | 0.00 |
| **$t_{50}(E\#L), s(E\#L)$** | **18** | **3046.87** | **0.00** | **1.00** | **1.00** | **3708.73** | **0.00** | **1.00** | **1.00** |

Models of snow crab larvae starvation survival ranked by Akiecke's Information Criterion with the best-fit model indicated by bold font. Model indicates the parameters in each model; parameters were linear function of factors where they are included parenthetically (see text for model details). Factors included pH treatment during embryo development (E) and during the larval experiments (L). K indicates the number of parameters in each model.

**Fig 1.**
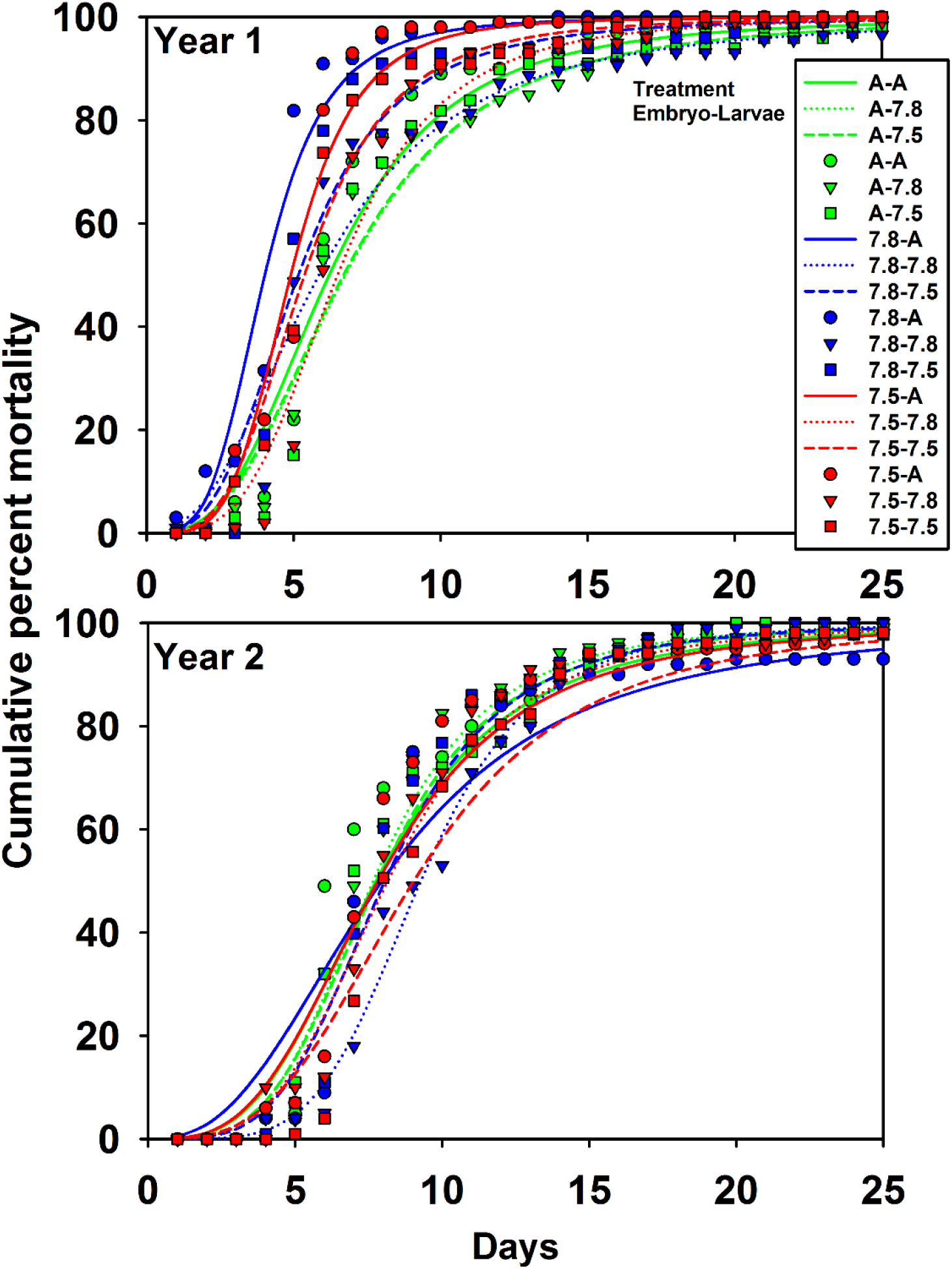
Effect of pH on larval snow crab starvation-survival. Cumulative percent mortality of snow crab larvae in starvation survival experiments in two years of experiments. Treatments represent the pH treatments during the embryo and larval portions of the experiments: A-Ambient pH, 7.8-pH 7.8, 7.5-pH 7.5. Points represent the average of five replicate trials and lines show the best fit curves (see Table 3 for parameter estimates).

**Table 3.** Snow crab larval starvation survival model parameters.

| Embryo | A | A | A | 7.8 | 7.8 | 7.8 | 7.5 | 7.5 | 7.5 |
| --- | --- | --- | --- | --- | --- | --- | --- | --- | --- |
| Larvae | A | 7.8 | 7.5 | A | 7.8 | 7.5 | A | 7.8 | 7.5 |
| Year 1 |  |  |  |  |  |  |  |  |  |
| $t_{50}$ | 6.20 | 6.75 | 6.69 | 4.02 | 5.86 | 5.15 | 4.90 | 6.49 | 5.36 |
| $s$ | -3.06 | -2.92 | -2.89 | -2.92 | -2.48 | -3.21 | -4.38 | -3.69 | -3.61 |
| Year 2 |  |  |  |  |  |  |  |  |  |
| $t_{50}$ | 7.75 | 7.65 | 7.81 | 7.95 | 9.27 | 8.05 | 7.82 | 8.15 | 9.03 |
| $s$ | -3.32 | -3.92 | -3.78 | -3.92 | -4.85 | -3.98 | -3.21 | -3.74 | -3.27 |

Parameter estimates for the best-fit model of snow crab larvae survival in starvation-survival experiments. Embryo and Larvae indicate the pH treatment at each stage. Parameters include the time to 50% mortality ( $t_{50}$ ) in days and a unitless slope parameter ( $s$ ). See text for model details.

In the first year, larval dry mass varied with both embryo and larval treatment and in the second year it varied only among embryo treatment (Fig. 2, Table 4). In the first year, larval mass was higher at hatching than after 7 days of holding but there was no difference among the three larval pH treatment. Additionally, larvae that hatched from embryos held at pH 7.5 were smaller than those in the other two treatments. In the second year larvae that hatched from embryos held at pH 7.8 were smaller than the other two treatments. Carbon and nitrogen content and the C:N ratio varied among both embryo and larval treatments and their interactions in both years (Fig 3, Table 4). Although the differences were statistically significant, there were no clear trends in the data either within or among years that showed consistent effects of either embryo or larval treatment and effect sizes were generally small (>5% differences). Calcium content varied among embryo and larval treatments in the first year but only among larval treatments in the second year (Fig. 4, Table 4). In the first year, there was a significant increase in calcium content between the larvae at hatching and after 7 days of holding with a trend for larvae held at pH 7.5 showing a non-significant increase. Further, larvae hatched from embryo held at pH 7.5 had lower calcium content than those from the other two treatments. In the second year, the only difference was a significant increase in calcium content between larvae at hatching and those held for 7 days; there were no differences among embryo or larval pH treatments. There were no significant differences in magnesium content among either embryo or larval treatments.

**Fig 2.**
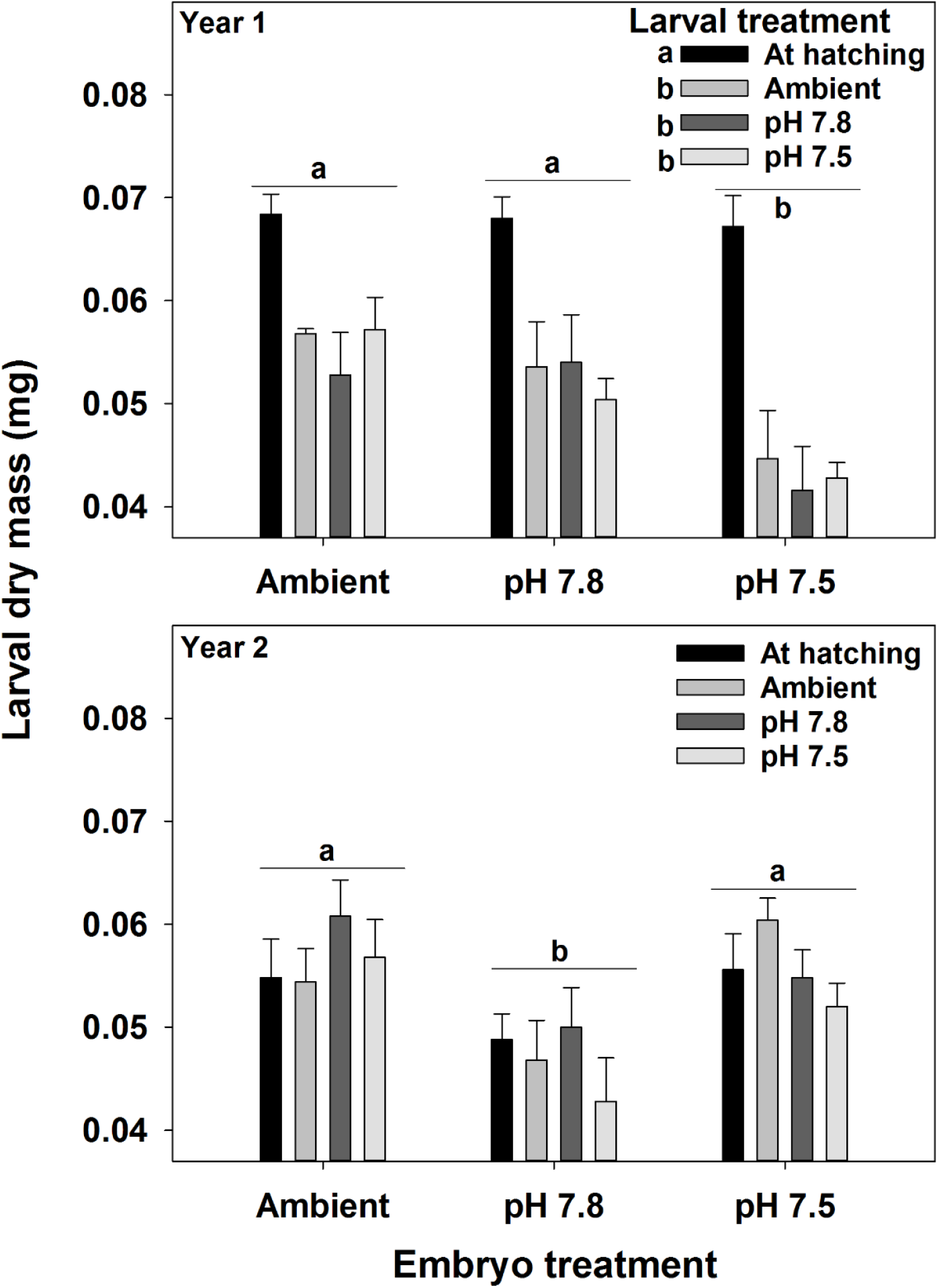
Water pH and snow crab larval mass. Effects of holding pH during the embryo and larval stages on the dry mass of snow crab larvae. Bars are means + 1 standard error. For larval treatments, “At hatching” represents larvae sampled immediately after hatching and Ambient, pH 7.8, and pH 7.5 represent larvae held at those pH treatments for 7 days. Statistically significant differences (Tukey’s test) among treatments are indicated with different letters.

**Fig 3.**
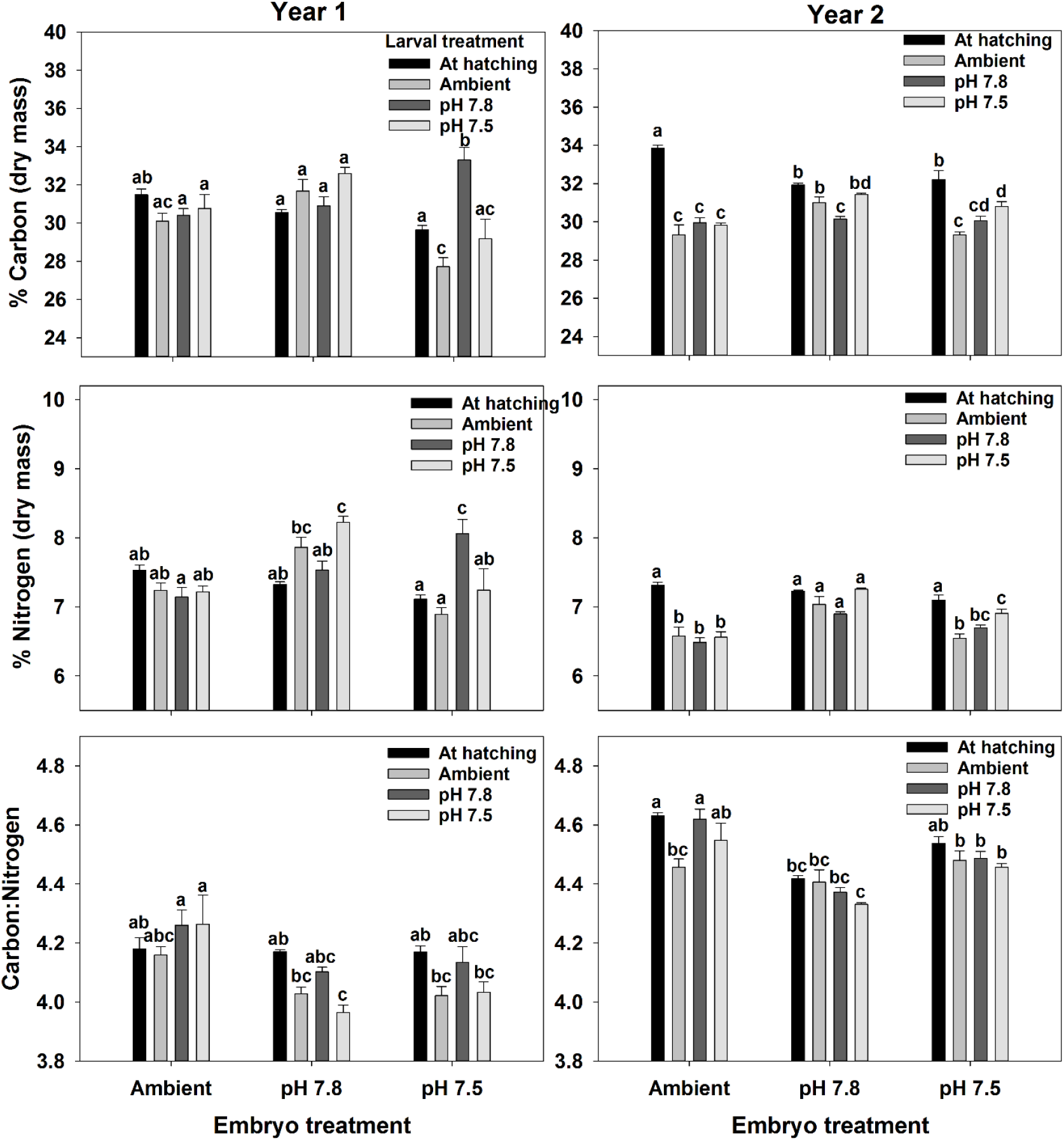
Water pH and snow crab larval elemental composition. Effects of holding pH during the embryo and larval stages on the carbon and nitrogen contents and the C:N ratio of snow crab larvae. Bars are means + 1 standard error. For larval treatments, “At hatching” represents larvae sampled immediately after hatching and Ambient, pH 7.8, and pH 7.5 represent larvae held at those pH treatments for 7 days. Statistically significant differences (Tukey’s test) among treatments are indicated with different letters.

**Fig 4.**
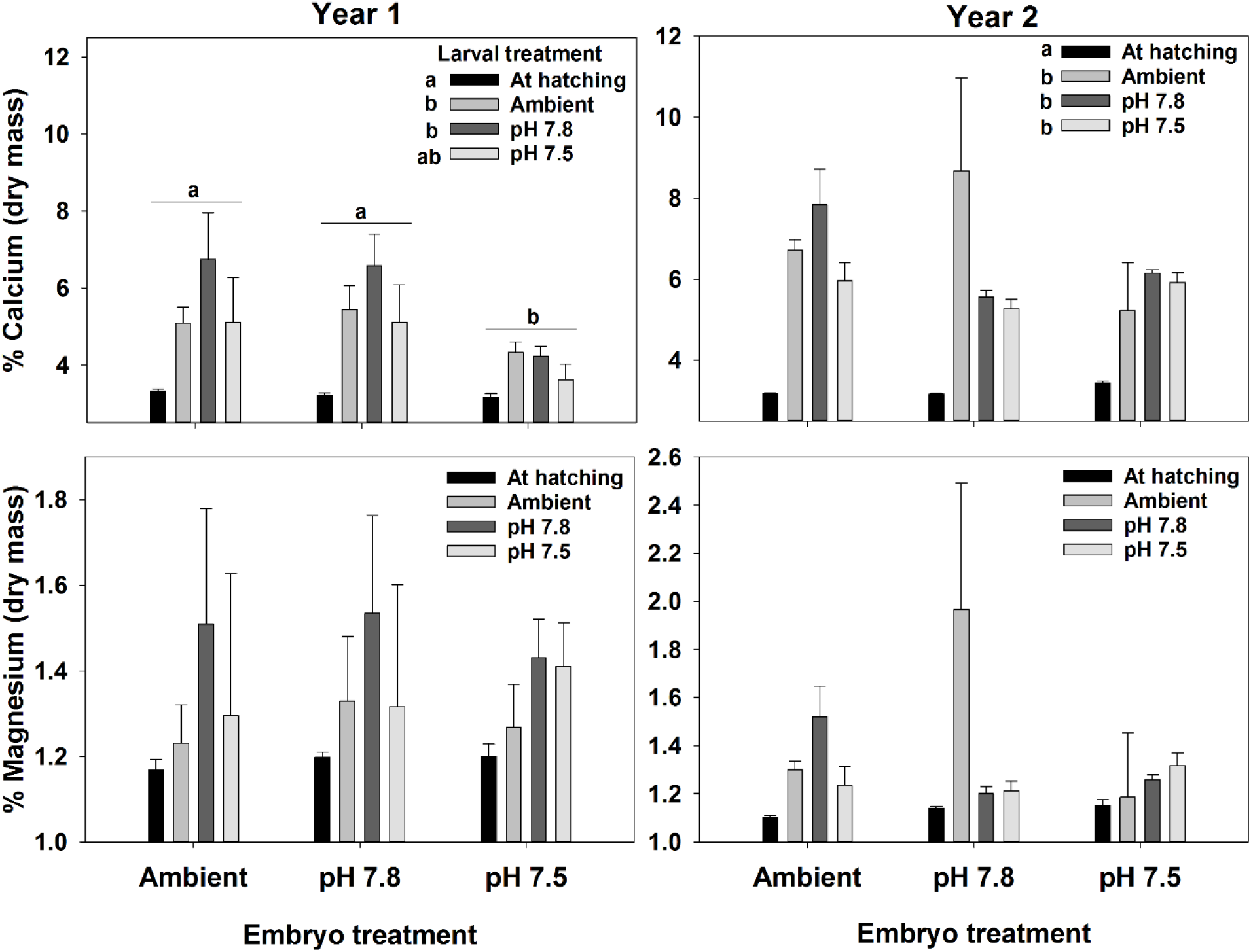
Water pH and snow crab larval calcium and magnesium content. Effects of holding pH during the embryo and larval stages on the calcium and magnesium contents of snow crab larvae. Bars are means + 1 standard error. For larval treatments, “At hatching” represents larvae sampled immediately after hatching and Ambient, pH 7.8, and pH 7.5 represent larvae held at those pH treatments for 7 days. Statistically significant differences (Tukey’s test) among treatments are indicated with different letters.

**Table 4.** Summary ANOVA results for snow crab larval experiments.

|  | Year 1 |  |  |  |  |  | Year 2 |  |  |  |  |  |
| --- | --- | --- | --- | --- | --- | --- | --- | --- | --- | --- | --- | --- |
|  | Embryo |  | Larval |  | E*L |  | Embryo |  | Larval |  | E*L |  |
|  | F | p | F | p | F | p | F | p | F | p | F | p |
| **C** | 7.547 | 0.001 | 5.233 | 0.003 | 8.829 | < 0.0005 | 3.804 | 0.029 | 61.894 | < 0.0005 | 9.948 | < 0.0005 |
| **N** | 12.854 | 0.000 | 3.030 | 0.038 | 9.309 | < 0.0005 | 31.240 | 0.000 | 36.009 | < 0.0005 | 6.929 | < 0.0005 |
| **C:N** | 13.729 | 0.000 | 5.497 | 0.003 | 2.605 | 0.029 | 37.567 | 0.000 | 5.417 | 0.003 | 2.386 | 0.043 |
| **Ca** | 4.500 | 0.016 | 7.619 | < 0.0005 | 0.735 | 0.624 | 0.855 | 0.432 | 12.031 | < 0.0005 | 2.000 | 0.084 |
| **Mg** | 0.028 | 0.973 | 1.148 | 0.340 | 0.115 | 0.994 | 0.730 | 0.487 | 2.061 | 0.118 | 1.962 | 0.090 |
| **Mass** | 9.291 | < 0.0005 | 22.689 | < 0.0005 | 1.145 | 0.352 | 9.995 | < 0.0005 | 1.038 | 0.384 | 0.860 | 0.531 |

Results of ANOVA for the effects of exposure to low pH at the embryo (E) and larval (L) stages on snow crab embryo in the first and second year of the experiments. Separate analyses were done for each year. Responses include carbon (C), nitrogen (N), calcium (Ca), and magnesium (Mg) contents; the C to N ratio; and the average larval dry mass (Mass).

Larval morphometry differed among treatments in both the first (Global R = 0.232, p < 0.0005) and second (Global R = 0.223, p < 0.0005) years (Fig. 5). Despite the statistical significance the low Global R value each year suggested actual differences among treatments were low. Simper analysis showed that the rostral-dorsal length and carapace width were the most important contributors to differences between treatment pairs in both years contributing to >50% of the difference between treatments in all cases (Table 5). In both years, larvae from embryos reared in Ambient water were in general slightly larger than those reared in lower pH waters in these metrics. However, effect sizes were small and smaller in the second than in the first year; effect sizes averaged 4.6% in the first year with a maximum of 13% whereas they averaged 3.4% in the second year with a maximum of 7.3%.

**Fig 5.**
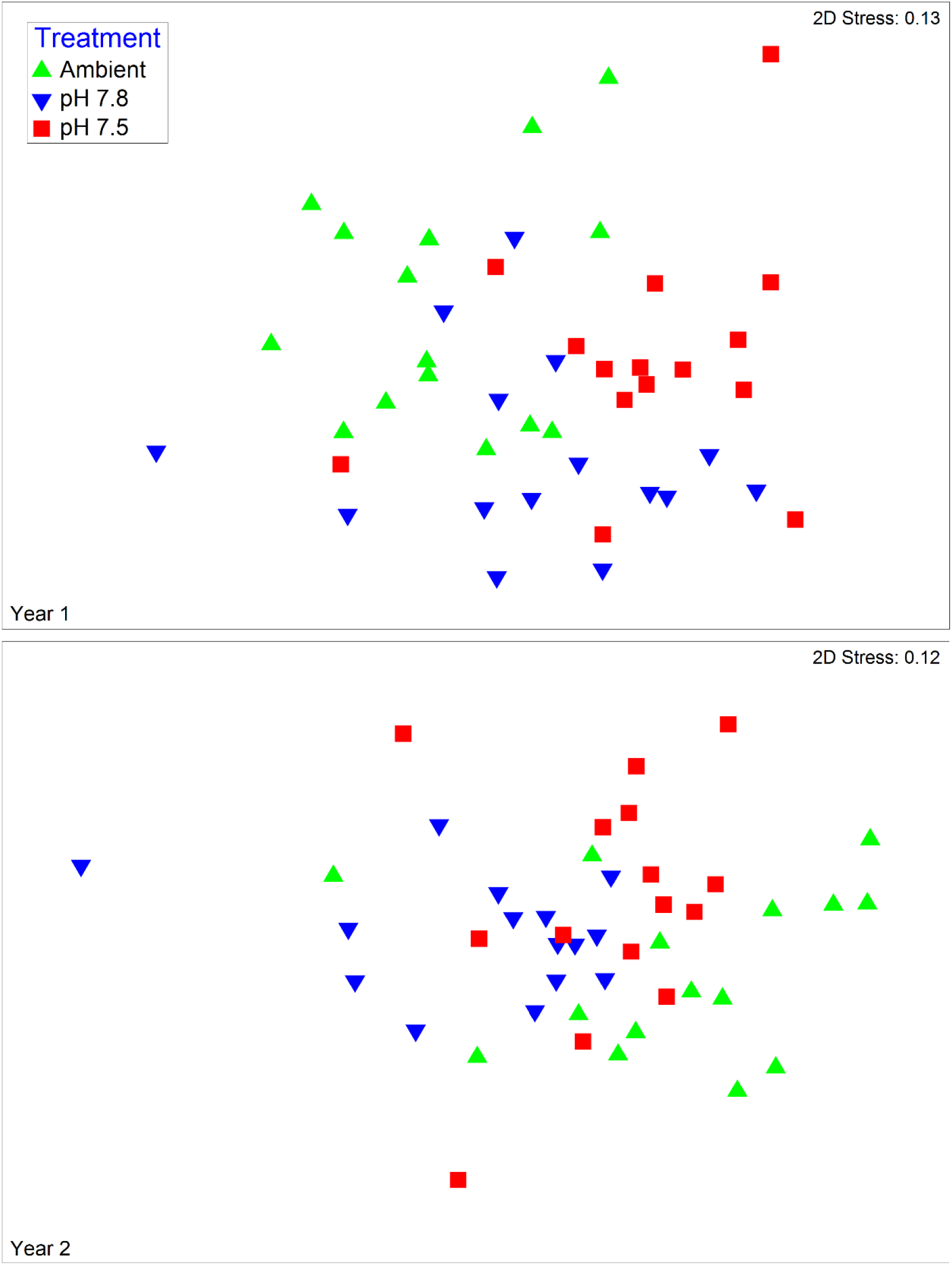
Effect of water pH on snow crab larvae morphometrics, Non-metric multidimensional scaling plots of morphometrics of snow crab larvae hatched from larvae held in three different pH waters in the first and second years of the experiment.

**Table 5.** SEMPER analysis results for effect of water pH on snow crab larval morphometry.

| Year 1 |  |  |  |  | Year 2 |  |  |  |  |
| --- | --- | --- | --- | --- | --- | --- | --- | --- | --- |
| Ambient vs. pH 7.8 |  |  |  |  |  |  |  |  |  |
| Variable | Mean (mm) |  | % Contribution | % Difference | Variable | Mean (mm) |  | % Contribution | % Difference |
|  | Ambient | pH 7.8 |  |  |  | Ambient | pH 7.8 |  |  |
| CW | 2.89 | 2.73 | 38.7 | 5.5 | RDL | 5.15 | 4.86 | 52.1 | 5.6 |
| PL | 1.68 | 1.83 | 20.3 | -8.9 | DSL | 2.18 | 2.07 | 14.9 | 5.0 |
| RDL | 5.24 | 5.2 | 14.0 | 0.8 | CW | 2.74 | 2.63 | 12.5 | 4.0 |
| LSL | 0.965 | 0.884 | 12.4 | 8.4 | RSL | 1.79 | 1.66 | 10.9 | 7.3 |
| DSL | 2.25 | 2.2 | 8.1 | 2.2 | PL | 1.78 | 1.71 | 6.2 | 3.9 |
| RSL | 1.67 | 1.74 | 6.6 | -4.2 | LSL | 0.91 | 0.905 | 3.5 | 0.5 |
| Ambient vs. pH 7.5 |  |  |  |  |  |  |  |  |  |
| Variable | Mean (mm) |  | % Contribution | % Difference | Variable | Mean (mm) |  | % Contribution | % Difference |
|  | Ambient | pH 7.5 |  |  |  | Ambient | pH 7.5 |  |  |
| CW | 2.89 | 2.68 | 34.5 | 7.3 | RDL | 5.15 | 5 | 35.9 | 2.9 |
| RDL | 5.24 | 5.07 | 24.5 | 3.2 | CW | 2.74 | 2.8 | 16.3 | -2.2 |
| LSL | 0.965 | 0.84 | 14.7 | 13.0 | DSL | 2.18 | 2.12 | 16.0 | 2.8 |
| PL | 1.68 | 1.76 | 11.2 | -4.8 | RSL | 1.79 | 1.66 | 14.9 | 7.3 |
| DSL | 2.25 | 2.14 | 9.3 | 4.9 | LSL | 0.91 | 0.923 | 9.7 | -1.4 |
| RSL | 1.67 | 1.67 | 5.8 | 0.0 | PL | 1.78 | 1.78 | 7.3 | 0.0 |
| pH 7.8 vs. pH 7.5 |  |  |  |  |  |  |  |  |  |
| Variable | Mean (mm) |  | % Contribution | % Difference | pH 7.8 | Mean (mm) |  | % Contribution | % Difference |
|  | pH 7.8 | pH 7.5 |  |  |  | Ambient | pH 7.5 |  |  |
| CW | 2.73 | 2.68 | 34.8 | 1.8 | RDL | 4.86 | 5 | 37.9 | -2.9 |
| RDL | 5.2 | 5.07 | 22.6 | 2.5 | CW | 2.63 | 2.8 | 26.0 | -6.5 |
| LSL | 0.884 | 0.84 | 13.8 | 5.0 | PL | 1.71 | 1.78 | 11.3 | -4.1 |
| PL | 1.83 | 1.76 | 10.5 | 3.8 | RSL | 1.66 | 1.66 | 9.8 | 0.0 |
| RSL | 1.74 | 1.67 | 9.3 | 4.0 | DSL | 2.07 | 2.12 | 8.4 | -2.4 |
| DSL | 2.2 | 2.14 | 9.0 | 2.7 | LSL | 0.905 | 0.923 | 6.6 | -2.0 |

Similarity percentage analysis of snow crab larvae morphometric measurements from larvae hatched from embryos held in water at three different pHs. Variables include carapace width (CW), lateral spine length (LSL), dorsal spine length (DSL), rostro-dorsal length (RDL), rostral spine length (RSL), and protopodite length (PL). Mean measurements for each treatment within each year are calculated. Variables are arranged by their percent contribution to differences between treatment comparisons. The % Difference represents the average percent difference between the two treatments with negative numbers indicating the second treatment is larger.

## Discussion

Snow crab larvae were very resilient to low pH conditions in these experiments suggesting that this life-history stage of this species may be well adapted to a broad range of pH conditions. In these experiments, we examined starvation survival and larval condition and were able to parse out direct effects from the carryover effects from oogenesis and embryogenesis. In general, effect sizes were small or non-significant and positive carryover effects from earlier life history stages reduced or eliminated the effects of OA suggesting the species is able to acclimate to large carbonate chemistry perturbations over long time periods. This in turn suggests that the stocks and associated fisheries may be less vulnerable to OA than that of other crab stocks.

There were no negative direct effects of exposure to low pH at the larval phase on snow crab larvae on any of the metrics measured here. This can be inferred by examining the differences (or lack thereof) among larval treatments within the embryos reared at ambient pH. Exposure to low pH as larvae did not affect mass, carbon or nitrogen content, or calcium or magnesium content. Further, the effect on starvation survival time was, if anything, slightly positive, increasing it by about half a day or around 10% in the first year, although not the second. This is consistent with the response of Tanner crab larvae in virtually identical experiments [33]. Effects of ocean acidification on larvae of crustaceans is mixed, there are many species whose larval phases are tolerant of low pH [45–47], but there are examples to the contrary [48–50]. It is not clear what factors are involved in determining the level of sensitivity to low pH; however, there is evidence that some species lack plasticity at the larval phase. Red king crab larvae, for example, have decreased survival and carbon content when held at low pH [49], but they do not alter their gene expression patterns [51]. Gene expression in snow crab larvae should be investigated to determine if they alter their physiology and, if so, what pathways they use.

Carryover effects from exposure to low pH at the embryo stage were generally larger than the direct effects. Such exposure reduced larval mass and size and substantially reduced starvation survival time by over a third in the first year. These results are also consistent with those of Tanner crabs where carryover effects from exposure at the embryo stage were larger than direct effects although effects sizes on snow crab were smaller [33]. Negative carryover effects, where exposure at one life history stage reduces performance at another stage is a common response [24,25,52]. In these cases, it is likely that increased energetic demands at these earlier stages, although they may not directly affect the earlier stage, affect processes later on.

However, snow crab larvae also experienced positive carryover effects. In the first year, the negative carryover effect on starvation survival was reduced or eliminated when the larvae were also held at low pH. This suggests that adaptive processes at the embryo stage altered the larval physiology in ways that adapted them to low pH. Further, positive carryover effects (based on a comparison of the results of the second year with those of the first) from oogenesis (also known as maternal effects) either eliminated or reduced, to the point where they were likely biologically insignificant, the negative carryover effects from embryo development on most of the parameters we measured including starvation survival, morphometry, and calcium content. And, beyond that, the carryover effects from oogenesis were additive to the adaptive processes that occurred during embryo development such that, in the second year, larvae that were hatched from females that had been held at low pH throughout egg and embryo development survived substantially longer than those held at ambient throughout.

It’s not clear what mechanism is involved in conferring the positive maternal effects. It could be selection, through selective mortality of females poorly adapted to low pH, or transgenerational plasticity [22]. Transgenerational and selective breeding experiments suggest that many species have the capacity to evolutionarily adapt to ocean acidification via natural selection [26, 53–56]. In the case of snow crabs, the mortality over the course of the experiment did not vary much between the treatments which makes this a less likely, though still possible, mechanism [57]. Alternatively, a stress response in the females could induce greater maternal investment in oocytes. The size and lipid content of crab eggs is a plastic response that can vary within and among populations and in response to environmental variables [58,59]; increased maternal investment can increase larval survival [60]. This is not well supported in the case of snow crabs as larval size (both in terms of mass and length) did not differ between embryos reared in ambient and pH 7.5 water in the second year and the carbon content was actually slightly lower in pH 7.5. Finally, epigenetic changes, particularly changes in DNA methylation can be induced by exposure to ocean acidification [61,62] and can be an adaptive response [63] and this possibility should be investigated further in snow crabs. Since these responses are not mutually exclusive any or all of them could be factors in this case.

In this study, we demonstrate that, at least for the parameters measured, snow crab larvae are resistant to low pH conditions and have the physiological plasticity to adapt to projected future carbonate chemistry conditions, especially since the embryos are similarly resistant [57]. However, more work needs to be done before it can be concluded the species as a whole will not be effected. First, resilience at one life history stage does not necessarily imply that for all other life history stages. Tanner crab larvae, for example, are very resistant to OA [33] whereas juveniles suffer increased mortality and decreased growth [34]. Likewise, European lobster, *Homarus gammarus*, larvae show very different sensitivities to ocean acidification when in different stages of larval development [64]. Thus experiments examining effects at other stages, particularly the post larvae and juvenile stages need to be performed. Further, ocean acidification will not occur in a vacuum and other stressors particularly increases in temperature and reductions in oxygen, are likely to co-occur; future work should then consider if these additional co-stressors will alter the response of snow crabs to reduced pH [65].

## Acknowledgments

We thank the RACE Groundfish and Shellfish Assessment Programs of the NOAA Fisheries Alaska Fisheries Science Center and the crews of the F/V Alaska Knight and F/V Arcturus for their assistance in securing crabs used in this study. We thank N. Gabriel, N. Sisson, A. Conrad (née Bateman), and staff of the Kodiak Laboratory for help performing the experiments. Previous versions of this paper were improved by comments from P. McElhany. The findings and conclusions in the paper are those of the authors and do not necessarily represent the views of the National Marine Fisheries Service, NOAA. Reference to trade names or commercial firms does not imply endorsement by the National Marine Fisheries Service, NOAA.

